# The SHR/Akr Y chromosome reveals repeated turnover of the rat pseudoautosomal region

**DOI:** 10.1101/2025.05.28.656580

**Authors:** Daniel W. Bellott, Helen Skaletsky, Jennifer F. Hughes, Laura G. Brown, Tatyana Pyntikova, Ting-Jan Cho, Natalia Koutseva, Sara Zaghlul, Dilziba Kizghin, Mayra Mendoza, Terje Raudsepp, Shannon Dugan, Ziad Khan, Qiaoyan Wang, Jennifer Watt, Kim C. Worley, Steven Scherer, Donna M. Muzny, Richard A. Gibbs, David C. Page

**Affiliations:** Whitehead Institute, Cambridge, Massachusetts 02142, USA; Howard Hughes Medical Institute, Whitehead Institute, Cambridge, Massachusetts 02142, USA; College of Veterinary Medicine and Biomedical Sciences, Texas A&M University, College Station, Texas 77843, USA; Human Genome Sequencing Center, Baylor College of Medicine, Houston, Texas 77030, USA; Department of Molecular and Human Genetics, Baylor College of Medicine, Houston Texas 77030, USA; Department of Biology, Massachusetts Institute of Technology, Cambridge, Massachusetts 02142, USA

## Abstract

Crossing-over between Chr X and Chr Y was first observed 90 years ago, in the brown rat, *Rattus norvegicus*. However, the sequence of the rat pseudoautosomal region (PAR) has remained a mystery. We produced a near-complete sequence of Chr Y from the SHR strain, along with nearly a megabase of sequence from both telomeres of Chr X. Both telomeric ends of Chr Y display extensive homology to Chr X, but no homology to the ancestral PAR of placental mammals. Using rat Y BACs probes for FISH on meiotic cells, we show that pairing almost always occurs between the tips of Yq and Xp, which are virtually identical in nucleotide sequence, but contain no protein-coding genes. Homology at the other ends of Chr X and Chr Y is likely the result of a recent transposition of five genes from Xq to Yp. These sequences are only 99.5% identical and pair infrequently, but show signs that it may have been pseudoautosomal in the ancestor of the rat genus. The rat Chr Y sequence presents opportunities for experimental studies of meiotic phenomena in a tractable model organism. The short PAR, with a correspondingly high recombination rate, represents a unique substrate for molecular studies of crossing-over. Likewise, the absence of extensively co-amplified testis gene families on the rat X and Y suggests they might serve as control for the intense competition between selfish elements that completely remodeled the mouse sex chromosomes.

## Introduction

The sex chromosomes of therian — placental and marsupial — mammals evolved from ordinary autosomes (Bellott et al. 2010, 2014; Lahn and Page 1999; Ohno 1967). After the advent of the testis-determining gene, *Sry* (Gubbay et al. 1990; Sinclair et al. 1990), on the nascent Chr Y, a series of events, most likely inversions on Chr Y, gradually suppressed X-Y crossing over (Bellott et al. 2014; Lahn and Page 1999). During meiosis, crossing-over physically links each pair of homologous chromosomes, providing the tension required to segregate them faithfully (Darlington 1937; McKim et al. 1993). On the sex chromosomes, crossing-over creates a pseudoautosomal region (PAR), where genes display partial sex linkage, and Chr X and Chr Y remain identical (Burgoyne 1982; Darlington et al. 1934; Koller and Darlington 1934). In the non-pseudoautosomal-Y (NPY) and non-pseudoautosomal-X (NPX), genes are completely sex-linked. Freed from the constraints imposed by crossing-over, the NPX and NPY have followed independent, and radically different, evolutionary trajectories.

In the absence of crossing-over, the NPY has been subject to genetic decay, and has lost most of its ancestral genes, while the NPX, which still engages in crossing-over during female meiosis, has retained almost all of them (Bellott et al. 2010, 2014; Ross et al. 2005; Skaletsky et al. 2003). The decay of NPY genes created a dosage imbalance between males and females. In therian mammals, this imbalance was resolved on a gene-by-gene basis (Jegalian and Page 1998; Naqvi et al. 2018; Ohno 1967; San Roman et al. 2023). In most cases, the upregulation of the NPX gene in both sexes, followed by epigenetic silencing of one of two NPX alleles in females, restores the ancestral gene dose, while creating two distinct epigenetic states for Chr X. In other cases, NPY gene loss was rescued by translocation of the NPY gene to an autosome (Hughes et al. 2015). The surviving ancestral genes on the NPY are enriched for broadly-expressed, dosage-sensitive regulators of key cellular processes that have NPX homologs expressed from each Chr X in females (Bellott et al. 2014; Lahn and Page 1997). These NPX-NPY gene pairs are predicted to maintain the ancestral gene dose and may play crucial roles in Turner syndrome as well as sex differences in health and disease.

A second consequence of the suppression of crossing-over has been the independent acquisition and amplification of multi-copy gene families predominantly or exclusively expressed in the germ cells of the testis by both the NPX and NPY (Hughes et al. 2010, 2012, 2020; Janečka et al. 2018; Mueller et al. 2013, 2008; Murphy et al. 2006; Paria et al. 2011; Ross et al. 2005; Skinner et al. 2015; Soh et al. 2014; Warburton et al. 2004). These gene families are embedded within ampliconic sequences — euchromatic repeats that display >99% identity over >10 kb. In some lineages, these ampliconic sequences completely dominate the euchromatic sequence of the NPY. In mouse and cattle, for example, amplicons make up 96% and 82% of the NPY, respectively (Hughes et al. 2020; Soh et al. 2014). Ampliconic structures are maintained by frequent gene conversion between paralogous repeats (Jackson et al. 2021a, 2021b; Rozen et al. 2003; Skaletsky et al. 2003; Swanepoel et al. 2020), but they are also prone to recurrent rearrangements that can dramatically remodel sex chromosome structure and lead to spermatogenic failure, sex reversal, or Turner syndrome (Lange et al. 2009; Repping et al. 2002, 2006).

Although crossing-over between Chr X and Chr Y was first reported ninety years ago in germ cells from the rat testis (Darlington et al. 1934; Koller and Darlington 1934), the sequence of the rat PAR remains unknown. Most placental mammals share an orthologous PAR (Raudsepp and Chowdhary 2016), but the extent of this region varies between species, depending on how many lineage-specific inversions occurred after divergence from the eutherian ancestor (Bellott et al. 2014; Raudsepp and Chowdhary 2016). While NPX genes are constrained to remain X-linked by the complexities of dosage compensation (Ohno 1967), PAR genes can tolerate translocation to autosomes, and this has been a frequent occurrence in the mouse (Blaschke and Rappold 1997; Disteche et al. 1992; Gianfrancesco et al. 2001; Milatovich et al. 1993; Palmer et al. 1995; Rugarli et al. 1995). At the extreme, this can result in the total loss of the PAR, as has occurred in marsupials (Sharp 1982; Solari and Bianchi 1975) and some rodent species (Borodin et al. 2012; Solari and Ashley 1977), where alternative mechanisms for sex chromosome segregation without crossing-over have evolved. Alternatively, the PAR can be extended by translocation from an autosome, as happened in the eutherian ancestor (Bellott et al. 2010; Graves and Watson 1991; Ross et al. 2005), and a second PAR can evolve through translocations between X and Y, like the human-specific PAR2 (Kvaløy et al. 1994).

On the rat sex chromosomes, crossing over occurs between the long arm of Chr Y and the short arm of the acrocentric Chr X (Joseph and Chandley 1984; Koller and Darlington 1934). It is therefore likely that the heterochromatic satellite arrays typically associated with centromeres and telomeres have thwarted efforts to assemble the rat PAR using clone-based and short-read sequencing technologies. Out of the 38 genes present on the ancestral placental PAR, 21 have orthologs identified in the rat genome, and all are either assigned to autosomes or the NPX (Maxeiner et al. 2020), suggesting that very little, if any, ancestral sequence remains in the PAR as a substrate for crossing-over.

The rat Chr Y was previously shown to encode members of the *Ssty* gene family (Ellis et al. 2011), a novel testis-specific gene family acquired in the rodent lineage. In the mouse, members of the *Ssty* family, along with other lineage-specific gene families, are extensively co-amplified on the NPX and NPY, as part of a hypothesized evolutionary arms race between the sex chromosomes, where Chr X and Chr Y compete to increase their probability of transmission to the next generation, starting during male meiosis (Burgoyne et al. 1992; Conway et al. 1994; Soh et al. 2014). Given the enormous difficulty of assembling the highly ampliconic mouse NPY, we targeted the spontaneously hypertensive rat (SHR) strain (Okamoto and Aoki 1963), which has the shortest Chr Y among common laboratory strains (B. Trask, personal communication). The spontaneously hypertensive phenotype in this strain has been mapped to Chr Y (Davidson et al. 1995; Ely et al. 1993; Ely and Turner 1990; Kren et al. 2001; Mattson et al. 2007), and because the effect of the SHR Y is mediated by androgen signaling (Ely et al. 1991, 1993), the multicopy *Sry* gene family has emerged as a leading candidate (Prokop et al. 2016, 2013; Turner et al. 2007, 2009).

Here we report a near-complete, telomere-to-telomere sequence of the rat (Rattus norvegicus, SHR/Akr strain) Chr Y, together with nearly a megabase of novel sequence from both telomeres of Chr X, including the PAR. In the rat lineage, none of the PAR genes from the placental ancestor remain; instead, repeated translocations between Chr X and Chr Y have established new regions of X-Y homology to stabilize sex chromosome segregation during male meiosis. The structure of the rat NPY most closely resembles that of mouse or bull, with a short arm containing ancestral X-Y gene pairs, and a long arm carrying newly acquired, lineage-specific amplified gene families expressed in the germ cells of the testis. However, compared to those species, rat Chr Y is relatively poor in genes and rich in satellite repeats. This may be because a dense array of the *Ssty* gene family has translocated to an autosome, potentially acting as an autosomal suppressor of sex-linked transmission ratio distortion, preventing selfish X-Y arms races from dominating the rat sex chromosomes. The combination of unusual features we observe on the sex chromosomes position the rat as a key model for advancing our understanding of the links between the evolution of the sex chromosomes and the mechanisms of meiosis.

## Results

### Sequencing, mapping, assembly and annotation of the SHR/Akr rat Y chromosome

We produced and assembled a near-complete sequence of rat Chr Y from the SHR/Akr strain. This assembly spans from telomere to telomere, including the pseudoautosomal region and centromere (Fig. 1a; Fig. S1; Data S1 & S2). To achieve a high level of contiguity and accuracy, we combined long-read whole genome shotgun sequencing data with the clone-based single-haplotype iterative mapping and sequencing (SHIMS) methodology (Bellott et al. 2018, 2022), which we previously used to sequence sex chromosomes in human, chimpanzee, rhesus macaque, mouse, bull, and chicken (Bellott et al. 2010, 2017; Hughes et al. 2010, 2012, 2020; Jackson et al. 2021a; Mueller et al. 2013; Skaletsky et al. 2003; Soh et al. 2014).

**Figure 1:**
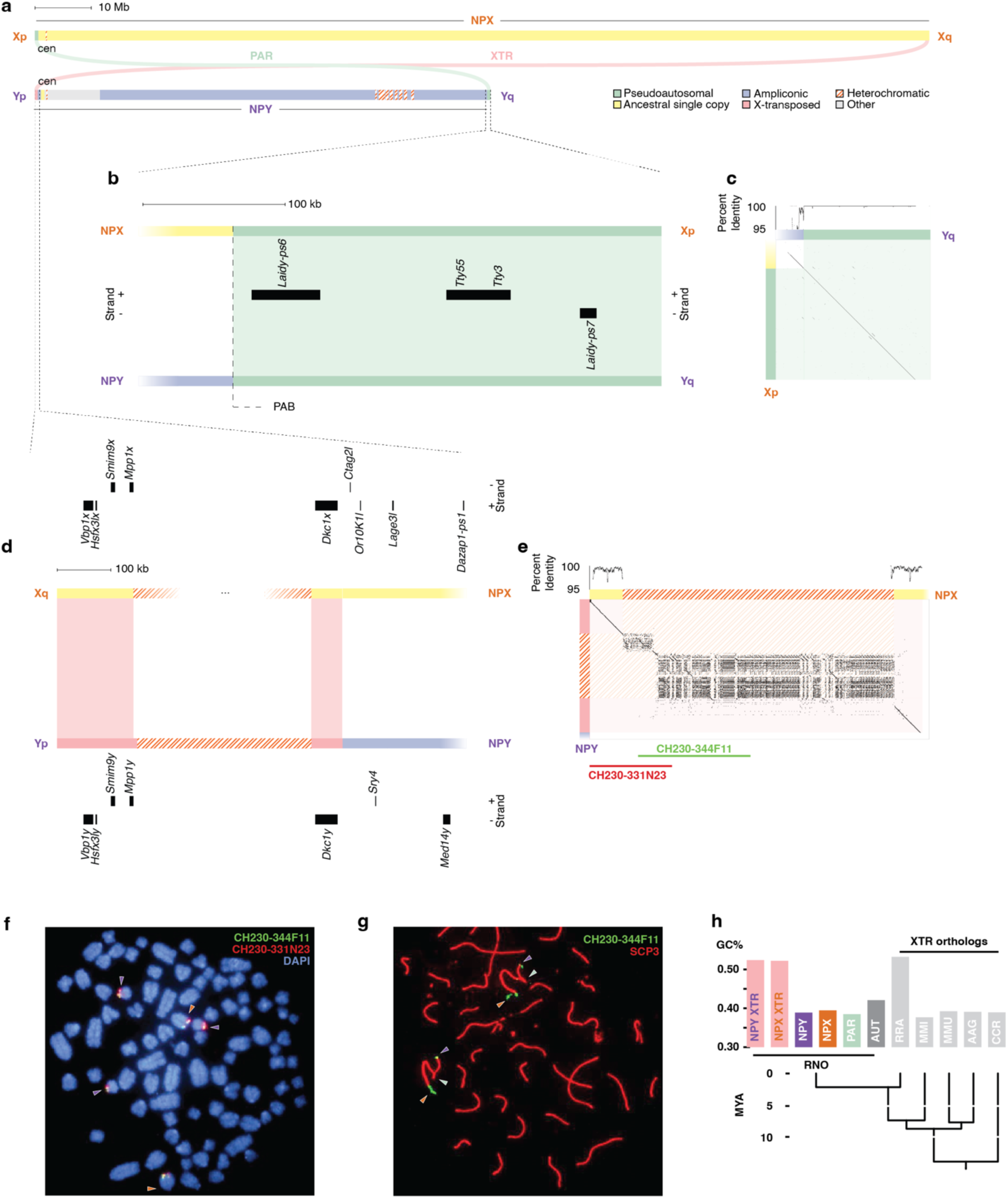
Structure of the rat sex chromosomes and pseudoautosomal region. (a) Schematic representation of rat Chr X and Chr Y, showing the whole of each chromosome, including the PAR. Shading indicates five classes of euchromatic sequence, as well as heterochromatic sequences. Colored arcs represent regions of >95% X-Y sequence identity. Scale bar, 10 Mb. (b) Distal 350 kb from the short arm of Chr X and long arm of Chr Y, enlarged to show PAR. Dashed line: PAB. Boxes, transcripts in PAR. Scale bar, 100 kb. (c) Square dot plot comparing the distal tip of the Y long arm to the X short arm. Each dot represents 100% identity in a 50-bp window. Above, graph of percent X-Y identity in 1-kb sliding window. (d) Distal 700kb from the long arm of the X and short arm of the Y, enlarged to show XTR. Boxes, transcripts on NPY and NPX. Scale bar, 100kb. (e) Square dot plot comparing the distal tip of the Y short arm to the X long arm. Each dot represents 100% identity in a 50-bp window. Above, graph of percent X-Y identity in 1-kb sliding window. (f) FISH with X-transposed region BAC probes CH230-344F11 (green) and CH230-331N23 (red) on mitotic metaphase chromosomes stained with DAPI (blue). Purple arrowhead, Chr Y; orange arrowhead, Chr X. (g) FISH with X-transposed region BAC probe CH230-344F11 (green) on meiotic prophase chromosomes of clone 9 (CRL-1439) cells, and anti-SCP3 antibody (red). Purple arrowhead, Chr Y; orange arrowhead, Chr X; green arrowhead, PAR. (h) Bar graph comparing GC content across regions of rat (RNO) genome against XTR orthologs in closely-related rodents: RRA, *Rattus rattus*; MMI, *Micromys minutis*; MMU, *Mus musculus*; AAG, *Apodemus agrarius*; CCR, *Cricetus cricetus*. Below, phylogenetic tree after Steppan and Schenk 2017 showing relationships between species.

By combining these complimentary approaches, we produced a model Chr Y assembly of 80 Mb (Methods; Data S1). Thirty contigs of Bacterial Artificial Chromosome (BAC) clones from a single SHR/Akr rat cover 70 Mb of the assembly; and within these contigs, our sequence is accurate to one nucleotide per megabase. To close the remaining 10 Mb of gaps resulting from unclonable heterochromatic satellite sequences, we used a combination of our own long reads from Oxford Nanopore Technologies and Pacific Biosciences Hifi reads from the closely-related SHRSP/BbbUtx strain (Kalbfleisch et al. 2023) to assemble BAC contigs into four scaffolds (Fig S1; Table S1; Data S2). The largest of these scaffolds spans 73 Mb, beginning at the telomere of the short arm of rat Chr Y, traversing the centromere and most of the long arm (Fig S1; Table S1; Data S2). At the other end, the long arm telomere is captured in a 2 Mb BAC contig that crosses the rat PAR and connects to the NPY (Fig. S1; Tables S1 & S2; Data S2). We used fluorescence in situ hybridization (FISH) and HiC mapping to order and orient scaffolds within the assembly (Figs S2 & S3; Methods). We estimate this model assembly is 99% complete.

To ensure that we captured any novel genes acquired by the rat sex chromosomes as a result of X-Y arms races similar to those we hypothesized in mice and cattle (Hughes et al. 2020; Soh et al. 2014), we generated voluminous transcriptomic data from the rat testis, generating 107 million 2x100 paired-end Illumina RNA-seq reads as well as 3.9 million full-length cDNA sequences with Pacific Biosciences Iso-seq (Methods). This permitted a comprehensive annotation of transcripts encoded by the rat Y (Tables S3 & S4; Data S3-S5), including 65 protein coding genes in 17 families and 95 non-coding transcripts in 40 families.

### Structure of the rat NPY

The rat Chr Y is a mosaic of sequence classes with distinct evolutionary histories (Fig. 1a). All sequences with homology to the ancestral autosomes are found within the 2 Mb short arm. At the tip of the short arm, the X-transposed sequences are the result of a recent X-to-Y transposition (Fig 1a; Table S2). This transposition restored ancestral sequences from the long arm of Chr X that had been lost from Chr Y before the divergence of placental mammals. Nearby, ancestral sequences have distant homologs on Chr X (Fig 1a; Table S2); these are the exceptional genes that survived from the ancestral autosome, despite widespread genetic decay on the rest of the NPY. The long arm is dominated by amplicons, euchromatic repeats that display greater than 99% sequence identity over more than 10 kb (Fig 1a; Table S2). At the distal tip of the long arm, a short pseudoautosomal region PAR (Fig. 1a; Table S2) crosses over with Chr X during meiosis, maintaining near-complete identity with allelic sequences on Chr X. Heterochromatic repeats are found at the centromere and near the telomere of the short arm, as well as an archipelago of heterochromatic islands spanning nearly 7 Mb in the middle of the long arm (Fig 1a; Table S2).

### Identification of the rat pseudoautosomal region

To identify the PARs of the rat sex chromosomes, we also generated targeted assemblies of each of the telomeres of Chr X (Fig 1, Table S1, Data S6 & S7). The sequences in these scaffolds are either absent or misplaced in previous assemblies of the rat genome.

The first of these two scaffolds spans 400 kb from the telomere on the short arm of Chr X and includes the PAR (Fig 1a & b). We observe that this Xp scaffold is virtually identical to the tip of Yq, differing by only five nucleotides over 282 kb (Fig 1c, Data S8). This is consistent with previous reports of pairing and synapsis between Xp and Yq during meiosis in male rats (Joseph and Chandley 1984; Koller and Darlington 1934), and with this sequence being pseudoautosomal. This sequence contains none of the ancestral PAR genes found across other eutherians (Raudsepp and Chowdhary 2016), but only carries three non-coding transcripts with homology to other transcripts in the NPY, including two pseudogenes from the *Laidy* gene family (Fig 1b, Table S3, Data S3). Like the *Ssty* family, *Laidy* has co-amplified NPX and NPY homologs in mouse (Arlt et al. 2020). In rat, as in mouse, only *Laidx* has coding potential (Arlt et al. 2020), and all *Laidy* copies are pseudogenes (Table S3, Data S3). We therefore conclude that this PAR is not ancestral, and arose by translocation of formerly NPY-specific sequence to Xp.

The second scaffold, from the long arm of Chr X, spans 515 kb from the telomere of the long arm of Chr X through a large bloc of heterochromatic repeats (Fig. 1a & d). This scaffold is 99.5% identical to the short arm of Chr Y over 207 kb of single-copy sequence, but contains approximately 4-fold more heterochromatin (Fig. 1e, Data S9 & S10). This region contains five X-Y gene pairs (Fig. 1d, Table 1). Four of these — *Dkc1x/y*, *Mpp1x/y Smim9x/y*, *Vbp1x/y* — are also located near the end of the long arm of the human Chr X, suggesting that this sequence transposed to the NPY from its ancestral region at the tip of the NPX relatively recently. Two BAC clones, CH230-344F11 and CH230-331N23 (Fig. 1e), produced probes specific to Chr X and Chr Y (Fig. 1f) in metaphase FISH. During prophase of male meiosis, CH230-344F11 hybridizes to the loose ends of the X and Y bivalent, opposite from the end where pairing occurs (Fig. 1g, Fig. S6), and synapsis of Xq and Yp is rare (< 2% of bivalents). Thus crossing over must primarily occur between Xp and Yq, and, given the level of sequence divergence, rarely, if ever, between Xq and the X-transposed sequence on Yp.

**Table 1:**
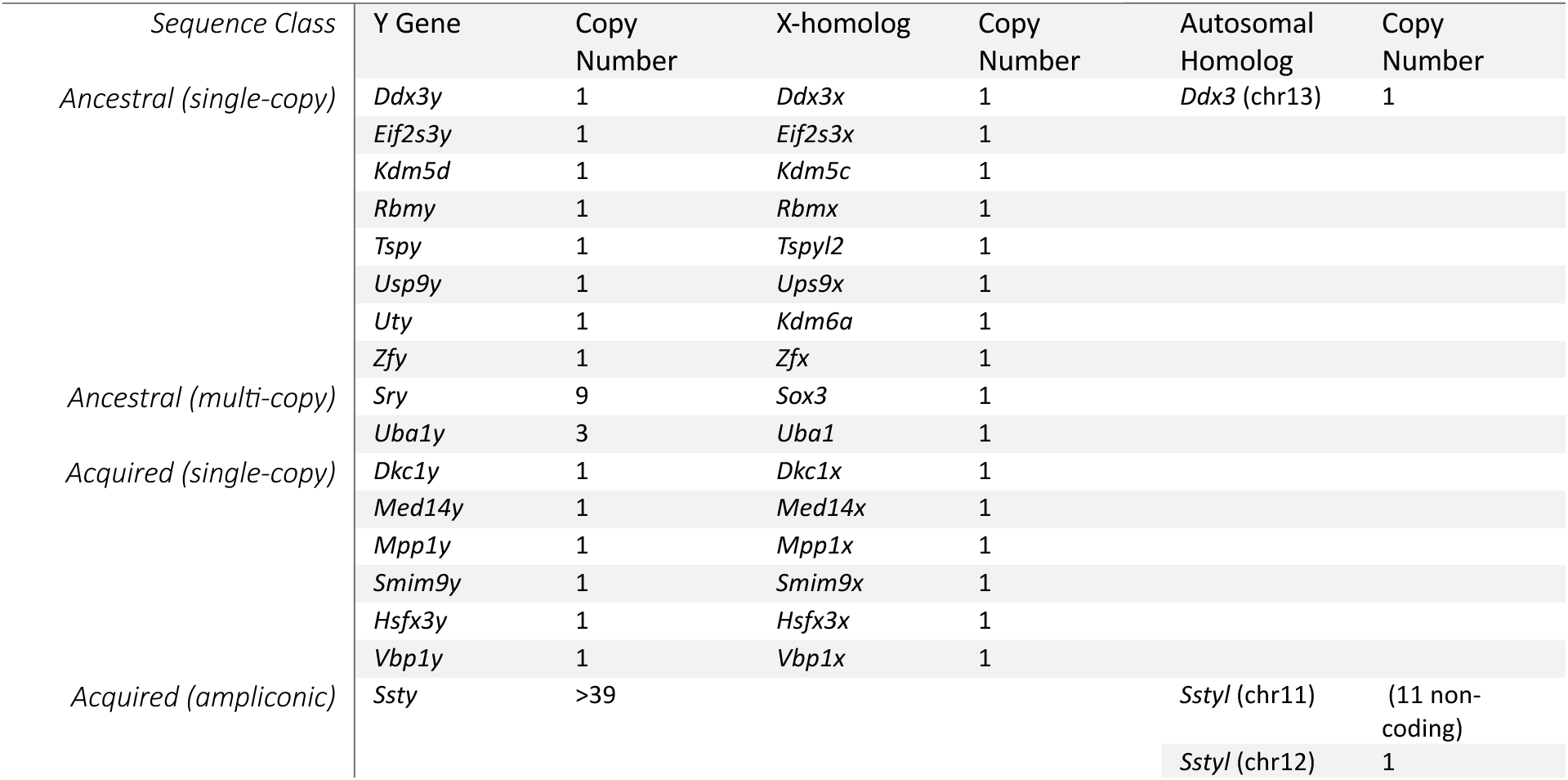
Rat Y genes and gene families and their X and autosomal homologs.

Although Y-linked XTR has diverged from related sequences on the NPX, there are signs that this region may once have engaged in crossing over. The GC content in the XTR is noticeably higher than in the rest of Chr Y (Fig. S1). We calculated that the GC content of the XTR and the paralogous sequence on the X is elevated compared to the remainder of the NPY and NPX, as well as the PAR and autosomes (Fig. 1h). The current PAR has GC content more typical of the rest of the NPY (Fig. 1h). One possible explanation is that the transposition that established the current PAR is extremely recent, and that the rat XTR formerly functioned as PAR; during that time, frequent crossovers, accompanied by GC-biased gene conversion, increased GC content in the XTR, but these processes have not yet had significant impact on the current PAR. Alternatively, the rat XTR may have originated from a sequence with ancestrally high GC content. We examined the XTR-orthologous regions on the NPX in several other rodents with highly-contiguous genome assemblies (*Cricetus cricetus*, *Mus musculus*, *Apodemus agrarius*, and *Micromys minutis*, and *Rattus rattus*) and found that all of them, except for the closely related black rat (Rattus rattus), have GC content that is more typical of NPX sequence (Fig. 1h). This suggests that this region transposed from the X to the Y and was used as a PAR after the divergence of *Micromys minutus*, but before the divergence from *Rattus rattus*, sometime in the last 7 million years (Steppan and Schenk 2017). Further investigation of rodent sequence orthologous to the rat PAR and XTR will clarify when these transpositions between the X and Y took place, and whether these sequences are pseudoautosomal in any other present-day species.

### Comparison with other placental Y chromosomes

The rat NPY (Fig 2a) most closely resembles the size and structure of the mouse (Fig. 2b) or bull NPYs. In each lineage, Chr Y is organized in a bipartite structure, with short arms bearing ancestral single-copy genes, and long arms with extensively amplified lineage-specific sequences (Hughes et al. 2020; Soh et al. 2014). We therefore compared the rat NPY sequence to that of the mouse (Fig. 2c, Fig S4). Despite the close evolutionary relationship between these two species (Fig. 2d), we found that very little sequence aligned outside of ancestral protein-coding genes (Fig. 2c, Fig S4). Even restricting our alignment to the conserved short arms (Fig 2c), where the mouse and rat share a near-identical set of ancestral genes (Bellott et al. 2014), very low stringency was required to detect sequence homology. This is likely a result of the rapid evolution of NPY sequence (Hughes et al. 2010), compounded by the accelerated rate of evolution in the rodent lineage (Li et al. 1996; Li and Wu 1987).

**Figure 2:**
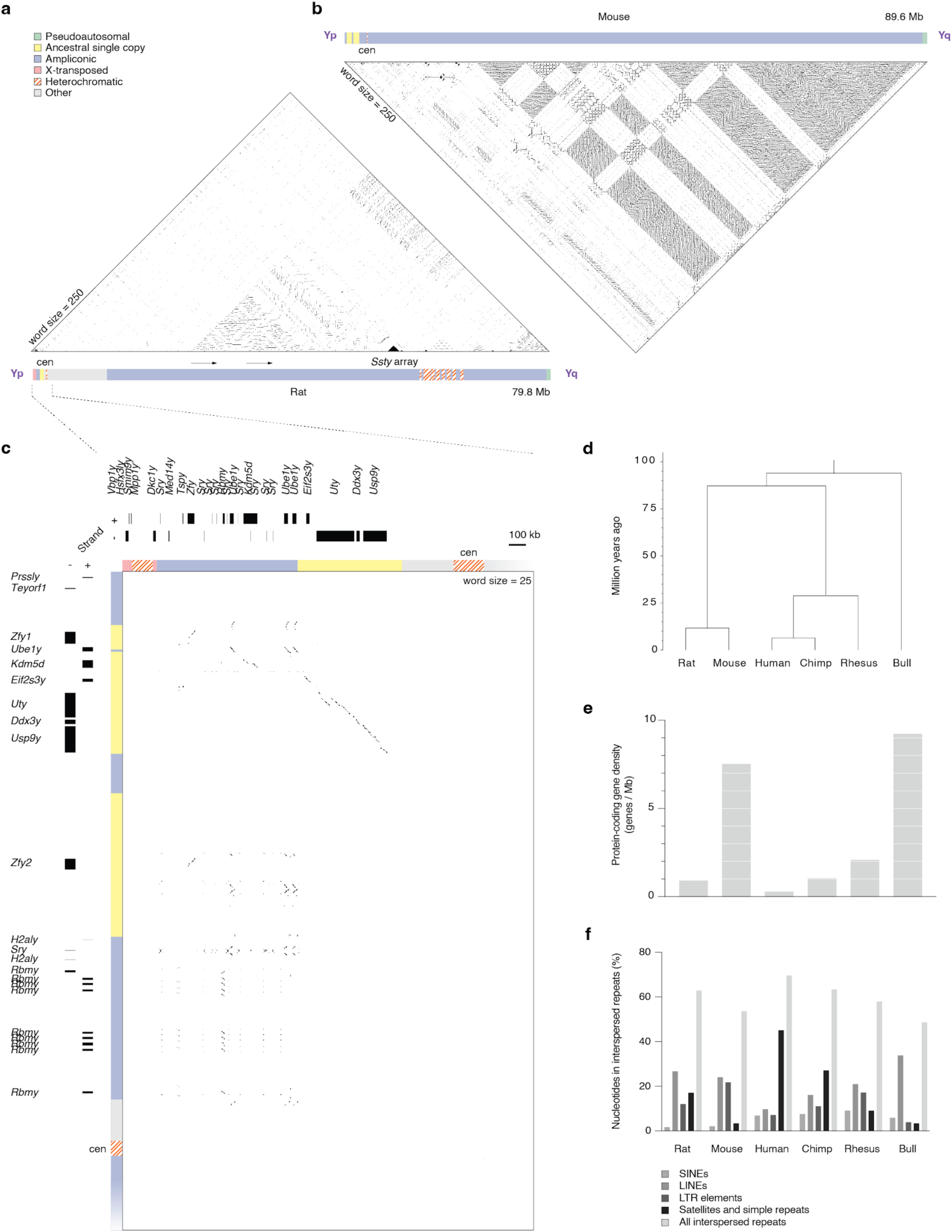
Comparison with other placental Y chromosomes. (a) Triangular dot plot. Each dot represents 100% identity in a 250-bp window between two sequences on the rat Y. Schematic representation of rat Chr Y, showing the whole of each chromosome, including the location of the largest amplicon and the main *Ssty* array. Shading indicates five classes of euchromatic sequence, as well as heterochromatic sequences. (b) Triangular dot plot and schematic of mouse Chr Y. (c) Square dot plot comparing the rat and mouse short arms. Each dot represents 100% identity in a 25-bp window. Boxes, transcripts on rat and mouse NPYs. Scale bar 100kb. (d) Phylogenetic tree of the relationships between analyzed Y chromosomes. (e) Bar chart of Chr Y gene density (genes / Mb) of the species in (d). (f) Grouped bar chart of interspersed repeat classes.

While members of the *Ssty* gene family are present on the long arms of both the rat and mouse NPYs, the ampliconic structures differ radically across species (Fig. 2a & b; Tables 1 & S3). On the long arm of the mouse NPY, *Ssty* (along with *Sly* and *Srsy*) is organized in a regular lattice of euchromatic ampliconic repeat units that make the long arm, and the entire mouse NPY, gene dense (Fig. 2b & e). The frequent gene conversion required to maintain the high identity between mouse NPY ampliconic sequences likely also purges interspersed repeats, resulting in interspersed repeat content typical of an autosome (Fig. 2f). In contrast, the amplicons of the rat NPY are more diverse and irregular (Fig. 2a). The largest ampliconic structure is a tandemly repeated sequence spanning 3.5 Mb at 99.5% identity, which contains no protein-coding genes (Fig 2a). *Ssty* copies are far more sparse than in mouse, with almost half concentrated in a compact tandem array spanning 2 Mb (Fig 1a). Combined with the acquisition of heterochromatic satellite arrays (Fig. 2a & f), this results in lower protein-coding gene density (Fig. 2e), along with higher interspersed repeat content (Fig. 2f) on the rat NPY. In these respects, the rat NPY more closely resembles those of primates (Fig. 2e & f), particularly the human NPY, which shares additional parallels. Like the rat NPY, the human NPY also recently acquired lineage-specific X-transposed regions and heterochromatic arrays (Skaletsky et al. 2003), although these are both evolutionarily younger and more massive.

### Structure of the short arm

The short arm of the rat NPY spans just over 2 Mb (Fig. 3a), and carries 26 protein-coding genes from 16 distinct families, all with homologs on Chr X (Fig. 3b; Table 1). Ten of these gene families are survivors from the ancestral autosomes (Table 1) (Bellott et al. 2014; Prokop et al. 2013). Two of those ancestral genes, *Sry* and *Ube1y*, are duplicated into multi-copy families within ampliconic structures (Fig. 3a & b; Table 1).

**Figure 3:**
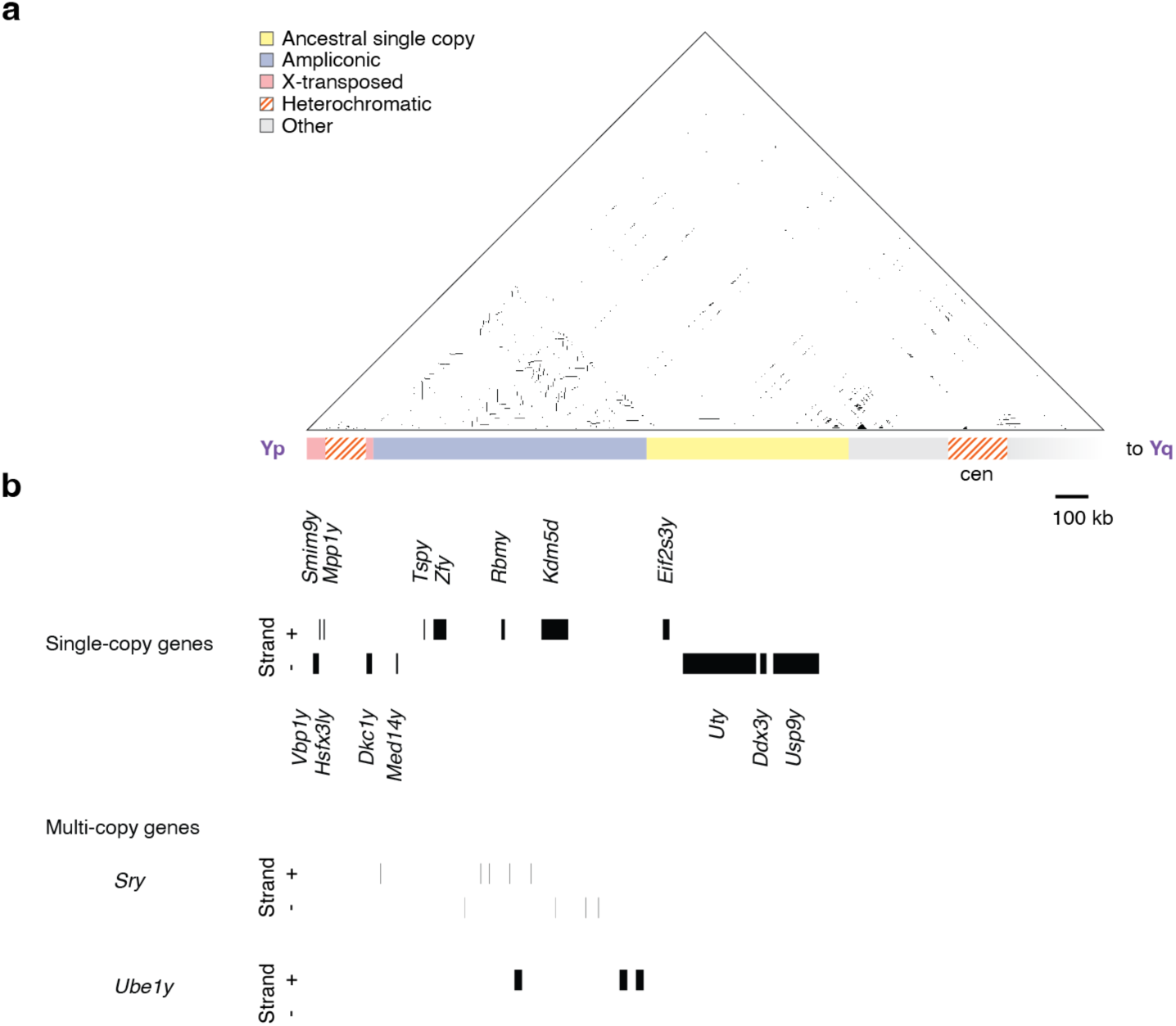
Structure of the rat Y short arm. (a) Triangular dot plot. Each dot represents 100% identity in a 250-bp window between two sequences on the rat Y. Schematic representation of the rat Y short arm. Shading indicates four classes of euchromatic sequence, as well as heterochromatic sequences. Scale bar 100kb. (b) Boxes, protein-coding transcripts on rat Y short arm.

The ten ancestral genes on the rat NPY were augmented by six genes recently acquired from Chr X by transpositions (Table 1). The intronless *Med14y* is the product of retrotransposition (Prokop et al. 2013), and is 93% identical to *Med14* on Chr X (Table S4; Data S9). The other five genes – *Vbp1y*, *Hsfx3ly*, *Smim9y*, *Mpp1y*, and *Dkc1y* – were transposed as a bloc from the distal tip of the long arm of Chr X (Figs. 1d & 3b; Table 1). These X-transposed genes on the NPY are about 99% identical to their NPX counterparts (Table S4; Data S8).

### The NPY exported *Ssty* genes to an autosome

In contrast to the short arm, none of the sequences on the long arm trace back to the autosomal ancestor of Chr X and Chr Y. Only a single protein-coding gene family is present on the long arm – the *Ssty* family (Fig 4a; Table 1). Of the 45 intact copies of *Ssty*, almost half are present in a single compact tandem array spanning 2 Mb (Figs. 2a & 4b). Each 85 kb repeat unit of this array carries a single copy of the *Ssty* gene (Fig. 4b). The remaining 24 protein-coding copies are scattered throughout the rest of the long arm among hundreds of non-coding transcripts (Fig. 4a; Table S3).

**Figure 4:**
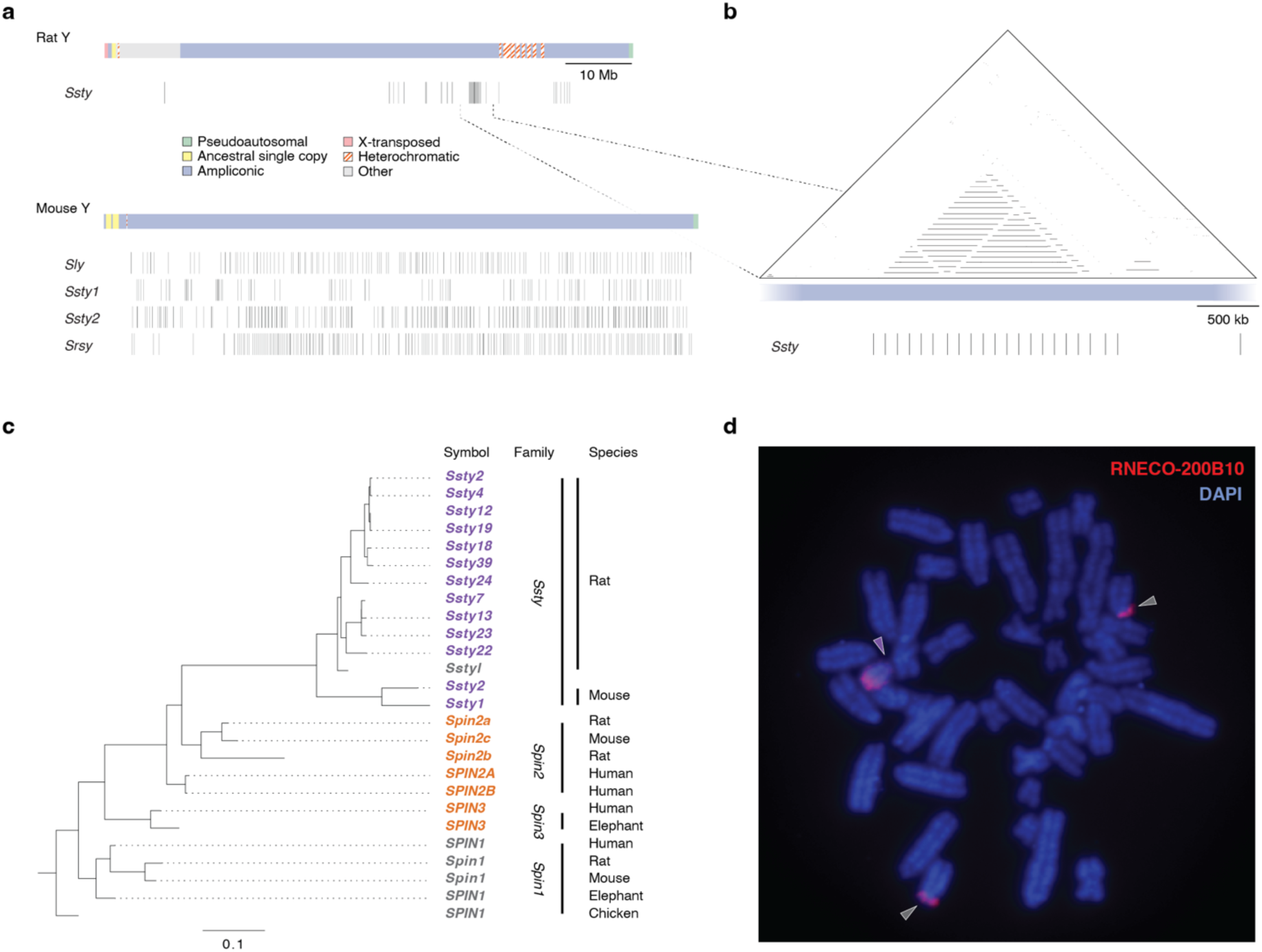
The ampliconic Ssty gene family. (a) Schematic representation of Chr Y from rat and mouse. Shading indicates five classes of euchromatic sequence, as well as heterochromatic sequences. Vertical bars indicate locations of ampliconic genes. Scale bar 10 Mb. (b) Triangular dot plot of the main rat *Ssty* array. Each dot represents 100% identity in a 250-bp window between two sequences on the rat Y. Scale bar 500kb. (c) Maximum-likelihood tree of selected *Ssty* family members and spindlin family homologs in placental mammals; gene names shaded by chromosomal location: purple, NPY; orange, NPX; grey, autosomal. (d) FISH with autosomal *Sstyl*-region BAC probe RNECO-200B10 (red) on mitotic metaphase chromosomes stained with DAPI (blue). Purple arrowhead, chrY; grey arrowhead, autosome.

The rodent-specific *Ssty* gene family is part of a larger spindlin family derived from the *Spin1* gene (Oh et al. 1997), which is located on Chr 17 in rat and is autosomal in other placental mammals (Fig. 4c). Before the divergence of placental mammals, a retrotransposition of *Spin1* to Chr X produced the intronless *Spin3*. Local duplications on Chr X created additional *Spin2* family members in the placental ancestor. Within the rodent lineage, *Spin2* was transposed to Chr Y, and amplified to form the *Ssty* family in the common ancestor of rat and mouse (Fig. 4a & c). In the rat, an array of *Ssty* genes have been transposed from Chr Y to an autosome (Fig. 4c & d). We identified an autosomal BAC probe containing *Ssty*-like sequences that hybridizes not only to the long arm of Chr Y, but also to the short arm of the smallest acrocentric autosome, chr11, at comparable intensity (Fig. 4d), suggesting that dozens of *Ssty*-like genes may be present in a large autosomal array. In our testis transcriptome data, we observe 12 distinct non-coding *Ssty*-like transcripts that do not originate from Chr Y and perfectly match existing female genome assemblies. The current rat reference assembly (GRCr8) assigns an array of *Ssty*-like genes to the short arm of Chr 3; but it is unclear whether this represents an artifact of the genome assembly process or a structural polymorphism among rat strains.

We previously observed that the mouse and bull NPYs are dominated by massively amplified testis-specific gene families, and that these ampliconic gene families have paralogs, amplified in parallel on the NPX (Hughes et al. 2020). We and others have hypothesized that these extensive co-amplification events are the result of recurrent evolutionary arms races between the sex chromosomes (Cocquet et al. 2012; Ellis et al. 2011; Hughes et al. 2020; Soh et al. 2014), where meiotic “driver” genes on the NPY act to increase the proportion of male offspring, and are opposed by “suppressors” on the NPX (Meiklejohn and Tao 2010; Presgraves 2008). In the mouse, the *Sly* and *Slx* gene families encode proteins that compete for binding to spindlin proteins, including those encoded by the *Ssty* family, and regulate thousands of genes during mouse spermatogenesis (Moretti et al. 2020). Manipulations that alter the dosage of *Slx* and *Sly* family members shift the sex ratio, with knockdowns and targeted deletions of *Slx* shifting the sex ratio in favor of males, while duplications of *Slx* or knockdowns of *Sly* shift the sex ratio in favor of females (Cocquet et al. 2009, 2012; Kruger et al. 2019). While studies of mouse collaborative cross lines did not observe a simple linear relationship between the *Slx*:*Sly* copy number ratio and sex ratio within each strain, several strains with biased sex ratios were found with extremes of *Ssty* copy number, suggesting that the effects of *Slx* and *Sly* copy number may be modulated by copy number variation among spindlin family members (Haines et al. 2021). With this in mind, we speculate that the colonization of a rat autosome by *Ssty* family members created an autosomal suppressor of both X- and Y-linked drive. The *Sstyl* array may have allowed the autosomes to adopt a potent armed neutrality and limit the scope of X-Y arms races in the rat, which could help explain why rat NPY ampliconic structures are more modest in scope than their counterparts in mouse.

### Gene Expression

We assessed the likely functions of NPY genes by measuring their expression patterns across a panel of nine adult rat tissues, including eight somatic tissues (Chang et al. 2023; Pattee et al. 2022; Schafer et al. 2015; Senko et al. 2022; van Heesch et al. 2019; Witte et al. 2021; Yang et al. 2020; Zeng et al. 2023) as well as the testis, and comparing their patterns with those of their NPX and autosomal homologs (Fig. 5a & b). We observed broad expression across adult tissues for the majority of single-copy NPY genes, including *Ddx3y*, *Eif2s3y*, *Kdm5d*, *Uty*, and all six single-copy acquired genes transposed from the NPX (Fig. 5a). These ten broadly expressed genes matched the expression patterns of their NPX homologs (Fig 5b). Both *Rbmy* and *Tspy* were already testis-specific in the common ancestor in the common ancestor of placental and marsupial mammals, and *Usp9y* and *Zfy* evolved testis-specific expression in the common ancestor of mouse and rat (Bellott et al. 2014), although all of their X-homologs are broadly expressed across somatic tissues (Fig 5a & b). Both ancestral multi-copy genes — *Sry* and *Uba1y* — are only very lowly expressed in adult tissues (Fig 5a), while *Ssty* is most highly expressed in the testis.

**Figure 5:**
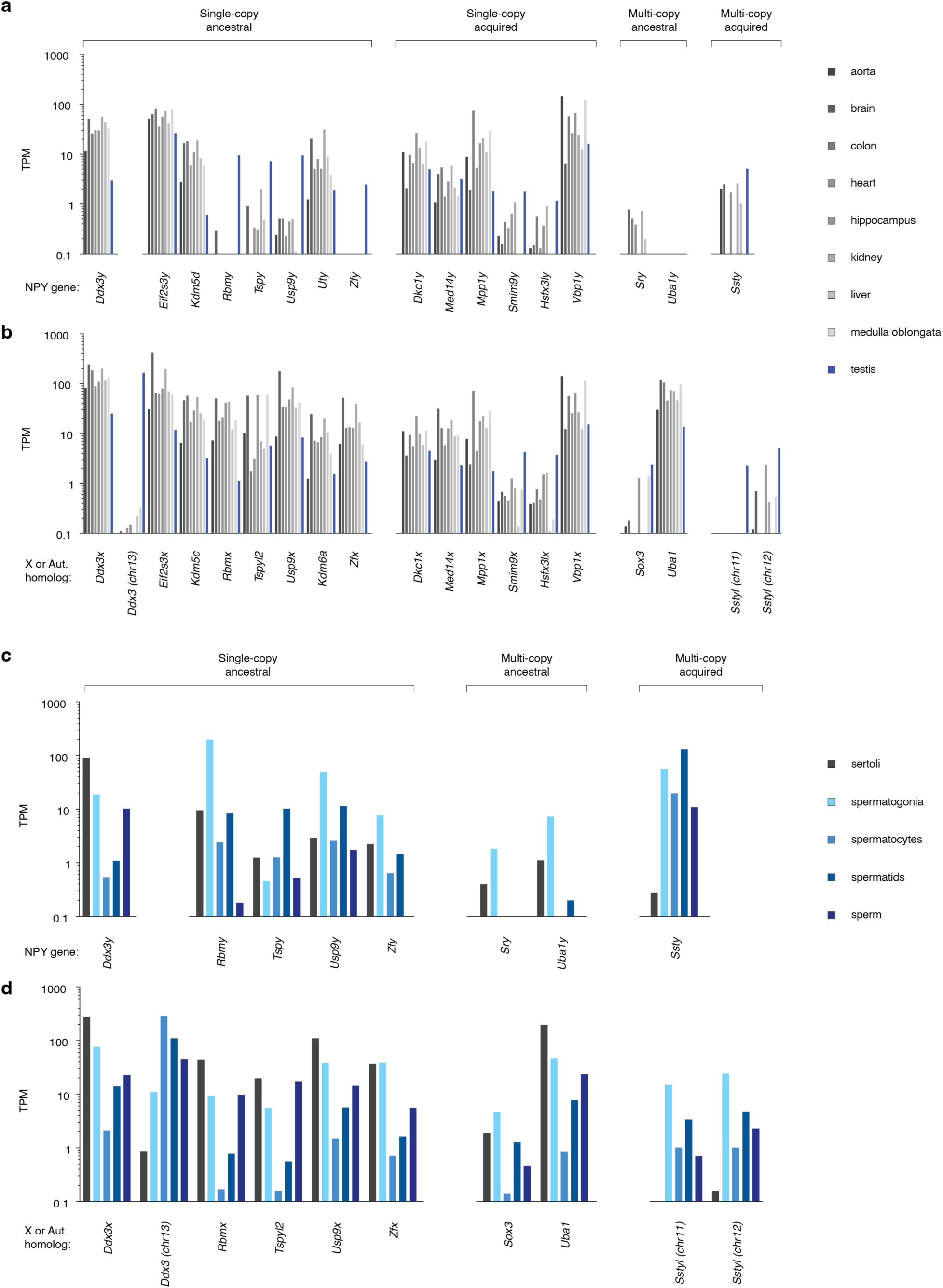
Expression of rat Y genes and their X and autosomal homologs. Expression of (a) Y genes and their (b) X and autosomal homologs in various adult tissues, as well as selected (c) Y genes and their (d) X and autosomal homologs in cell fractions of adult testes, as measured by RNA sequencing. For multi-copy and ampliconic genes, expression is the sum across all copies.

To further investigate the functions of NPY genes during male germ cell development, we examined expression across germ cell fractions from the testis (Chalmel et al. 2014; Fice and Robaire 2023; Li et al. 2025; Wang et al. 2022), including mitotic spermatogonia, meiotic spermatocytes, post-meiotic spermatids, and mature sperm, along with somatic Sertoli cells (Fig 5c & d). Among the ancestral genes with strongest expression in testis, *Rbmy*, *Usp9y*, and *Zfy* are predominantly expressed in germ cells, with a peak in spermatogonia, suggesting a role in maintaining or amplifying the pool of male germline stem cells (Fig. 5c). In contrast, *Tspy* is most highly expressed in post-meiotic cells (Fig. 5c), consistent with previous observations that rat *Tspy* has diverged from the expression pattern observed in other mammals and is expressed in elongating spermatids, where its protein product is hypothesized to act as a histone chaperone (Kido and Lau 2006).

Most broadly-expressed NPY genes have low expression in adult testis (Fig 5a), and within the testis, all NPY and NPX genes show a dip in spermatocyte expression (Fig. 5c & d), a consequence of meiotic sex chromosome inactivation (MSCI) (McKee and Handel 1993; Monesi 1965). *Ddx3x* and *Ddx3y* are extremely dosage sensitive, governed by elaborate cross- and auto-feedback regulation to maintain the correct combined *Ddx3x* and *Ddx3y* transcript levels (Rengarajan et al. 2025). *Ddx3* is an autosomal retrogene derived from *Ddx3x* in the ancestor of rodents, with strong testis-specific expression (Fig. 5b); within the testis, *Ddx3* has its highest expression in spermatocytes (Fig. 5d), when *Ddx3x* and *Ddx3y* levels are lowest due to MSCI (Fig. 5c & d). This strongly suggests that *Ddx3* evolved to compensate for the low combined dose of *Ddx3x* and *Ddx3y* during meiosis. We note that the mouse ortholog of *Ddx3*, *D1Pas1*, is required for the completion of meiotic prophase in male germ cells (Inoue et al. 2016).

The *Ssty* family and its autosomal paralogs are only lowly expressed outside testis (Fig. 5a & b); within the testis, they appear to be germ cell-specific, with the highest expression in post-meiotic spermatids (Fig 5c & d). This expression pattern is consistent with what is observed in mouse (Comptour et al. 2014; Soh et al. 2014; Touré et al. 2004), suggesting that *Ssty* genes may play similar roles in transmission distortion in both species, and supporting the idea that autosomal *Sstyl* genes could act as suppressors of *Ssty* drive.

## Discussion

Using a combination of SHIMS and long-read whole genome shotgun sequencing, we have assembled the Chr Y sequence of the SHR/Akr rat strain. Rat and mouse Chr Y are organized similarly, with a short arm carrying ancestral genes, and an ampliconic long arm with acquired sequences. But, despite this overarching similarity and their close evolutionary relationship, rat Chr Y has rapidly diverged from that of the mouse. While both mouse and rat feature ampliconic sequences containing members of the *Ssty* gene family, the mouse has acquired novel genes families in hundreds of copies and its Y arm is extremely structurally complex; in contrast, amplification on the rat Y is subdued; almost half of *Ssty* gene family members are confined to a narrow tandem array on the long arm, while others have decamped from Chr Y for autosomes. Even on the short arm, differential amplification and gene loss led to *Sry* and *Ube1y* becoming amplified in rat instead of *Rbmy* and *Zfy* in the mouse, and the ancestral *Prssly* gene relocated to an autosome in the rat. While ancestral gene survival and the acquisition and amplification of novel genes have been major themes in the evolution of other eutherian Y chromosomes, the trajectory followed by rat Chr Y has involved a remarkable amount of gene export to the autosomes.

The sequence of the rat PAR presents unparalleled opportunities for experimental studies of meiotic recombination in a mammalian model organism. During every male meiosis, an obligatory crossover occurs within a PAR spanning only 300 kb – this creates an extremely high recombination rate of approximately 166 centimorgans per megabase. A recombination rate orders of magnitude higher than the genome average might be expected to have profound effects on sequence evolution within the PAR. However, we observe that the PAR does not differ dramatically from the composition of the rest of the rat Y with respect to the abundance of interspersed repeats or GC content. This implies that the rat PAR originated very recently. Future studies of additional rat strains and closely related species within will reveal whether the current rat PAR is strain-specific, common to all rats, or shared with closely related species. The X-transposed region, with its noticeably elevated GC content, may have once functioned as a PAR in the past. Neither sequence is homologous to the ancestral eutherian PAR, suggesting that the PAR has turned over at least twice in the rat lineage after the divergence from mouse.

These turnover events underscore the dynamic nature of PARs and their potential to cause rapid structural change on the sex chromosomes. Transpositions between Chr X and Chr Y generate novel PARs (Kvaløy et al. 1994), and transpositions from autosomes to the PAR can extend them(Murata et al. 2012; Watson et al. 1990), while the PAR can shrink either because of inversions at the PAB that extend the NPX and NPY at the expense of the PAR (Lahn and Page 1997), or though transpositions from the PAR to the autosomes (Blaschke and Rappold 1997; Disteche et al. 1992; Gianfrancesco et al. 2001; Milatovich et al. 1993; Palmer et al. 1995; Rugarli et al. 1995). The meiotic products of unequal crossing over between the PAR and NPX (or NPY) will include both extended and deleted PARs (Mensah et al. 2014; Poriswanish et al. 2025). Like other regions of high recombination, PARs may be subject to the ‘hotspot paradox’ (Boulton et al. 1997; Coop and Myers 2007) – that recombinational mechanisms systematically disrupt ‘hotter’ alleles that initiate crossovers and replace them with their ‘colder’ homologs. This creates a selection pressure for structural variants that disrupt the recombinational potential of a PAR that only intensifies as a PAR grows shorter. We speculate that PARs are metastable; that as long as the drive to generate colder PARs is balanced by selection to maintain crossing-over, the PAR can persist, but that addition of a second PAR can destabilize the first, leading to turnover. Achiasmate mechanisms for the segregation of the sex chromosomes, of the type that have evolved repeatedly among rodents (Borodin et al. 2012), may provide an escape from the instability caused by repeated rounds of PAR turnover. Future cross-species comparisons will reveal the evolutionary history, genomic mechanisms and meiotic consequences of PAR turnover.

As the leading mammalian model organisms, the stark contrast between the ampliconic structures of the mouse and rat sex chromosomes will facilitate experimental tests of hypothesized sex chromosome meiotic drive. Targeted deletions and knockdowns of rat *Ssty* family members on Chr Y and autosomes, as well as related spindlins on Chr X will reveal their effects on the sex ratio and male fertility. In vitro binding assays, similar to those recently employed on mouse spindlin family members (Arlt et al. 2025), will identify the specificity of rat SSTY proteins and validate their ability to interact with SYCP3 family members and compete with each other. It will also be important to examine the roles of the abundant non-coding transcripts on the rat Y, some of which (e.g. *Laidy-ps*) are related to suspected drive elements that retain coding potential in the mouse. Comparisons between mouse and rat will reveal whether meiotic drive is due to the encoded mouse proteins, some function of the transcripts shared between species, or whether their promoters act as suppressors of meiotic drive by acting as a sink for SYCP3/SPIN complexes.

Lastly, sequence of a Chr Y from the SHR/Akr rat strain will serve as an important reference for genotype-phenotype correlations across rat strains. The SHR strain was developed by selectively breeding rats from the Wistar Kyoto (WKY) strain for higher blood pressure (Okamoto and Aoki 1963) and this phenotype was subsequently mapped to the SHR Chr Y (Ely et al. 1993). Because the effect of the SHR Y is mediated by androgen signaling (Ely et al. 1991, 1993), efforts to identify the Y-linked gene responsible for the SHR phenotype have focused on *Sry* (Prokop et al. 2016, 2013; Turner et al. 2007, 2009). Future comparisons of Chr Y between SHR/Akr and related normotensive strains may reveal additional differences in the complement of *Sry* family members, or differences in sequence or copy number elsewhere on Chr Y, that contribute to the spontaneously hypertensive phenotype. Large-scale structural variation in Chr Y has long been documented across various rat strains (Hungerford and Nowell 1963; Kodama and Sasaki 1977; Zieverink and Moloney 1965), and the ampliconic nature of rat Chr Y makes it vulnerable to genomic rearrangements. The reference of the SHR/Akr Chr Y will be essential for designing targeted genetic manipulations of Chr Y to test hypotheses about the genetic basis of the spontaneously hypertensive phenotype. Unravelling the mechanisms that connect Chr Y to high blood pressure in rat will have profound implications for understanding sex differences in human health and disease.

## Methods

### BAC selection and sequencing

We sequenced 1004 clones (Table S1) from two custom BAC libraries constructed by Amplicon Express using the SHIMS strategy (Bellott et al. 2018, 2022; Kuroda-Kawaguchi et al. 2001). Both libraries are derived from a single male donor from the SHR/Akr rat strain (Charles River Laboratories). We estimated the error rate in finished sequence by counting mismatches in alignments between overlapping clones.

### Long-read sequencing

To close gaps between BAC contigs, we generated additional long-read sequence data from an SHR/Akr male (SRR32833872) and an SHR/NCrl female (SRR32834695) rat. Briefly, we pulverized liver tissue with a mortar and pestle under liquid nitrogen and extracted high molecular weight DNA from the powdered tissue using the Monarch HMW DNA Extraction Kit for Tissue (New England Biolabs #T3060L). We prepared sequencing libraries with the Ultra-Long DNA Sequencing Kit V14 (Oxford Nanopore Technologies SQK-ULK114). We sequenced libraries on the Oxford Nanopore Technologies PromethION 2 Solo using R10.4.1 flow cells and used the Flow Cell Wash Kit (Oxford Nanopore Technologies EXP-WSH004-XL) to unblock pores and reload libraries twice during a 72-hour run. We used electronic subtraction to identify NPY-specific reads by comparing sequences from the male and female rats. All connections between BAC contigs are supported by nanopore reads from the SHR/Akr male rat; we used PacBio HiFi reads from the closely related SHRSP/BbbUtx strain (SRR17888679) (Kalbfleisch et al. 2023) to error-correct these contigs and extend into heterochromatic regions (Data S11).

### Fluorescence in situ hybridization (FISH)

We performed fluorescence in situ hybridization (FISH) using sequenced BACs as probes for rat Chr Y. We derived rat embryonic fibroblasts from a male of the SHR/CRL strain (WHT5890), and a female of the WKY strain (WHT5873), and obtained tetraploid liver epithelial cell line clone 9 (CRL-1439) from ATCC. We performed metaphase FISH as previously described (Saxena et al. 1996). We performed synaptonemal complex surface spreads on rat testes using rabbit anti SCP3 primary antibody (NB300-232, Novus Biologicals), with donkey anti-rabbit Alexa Fluor594 as a secondary antibody (ab150105, Abcam), and biotin-labeled BACs detected with streptavidin-Alexa Fluor488 (S11223, Invitrogen), following published protocols (Ma et al. 2006; Peters et al. 1997). FISH followed by immunostaining and immunostaining followed by FISH produced identical results.

### HiC-mapping

To confirm the order and orientation of sequence contigs within scaffolds, we mapped short-read data from the closely related SHRSP/BbbUtx strain (SRR17888680) (Kalbfleisch et al. 2023)to our Chr Y sequence, along with Chr X and autosomes from the GRCr8 (GCF_036323735.1) assembly, using minimap2 (version 2.26-r1175) (Li 2018). We used glistmaker (version 4.0) (Kaplinski et al. 2015) to identify all genome-wide unique 32-mers in our assembly, and filtered our HiC reads to retain only mate pairs overlapping these positions. For each pair of 50-kb windows at the flanks of sequence contigs in our assembly, we counted the number of mate pairs connecting the two flanks.

### Sequence annotation

We electronically identified interspersed repeats using RepeatMasker (Smit et al. 2020). To annotate transcripts, we generated and sequenced libraries for the Illumina HiSeq 2000 (SRR1001913, 107 million spots), and the PacBio Sequel II (SRR26701686, 3.9 million spots) from the adult testis of an SHR/Akr strain rat (Table S). We identified protein coding-genes as previously described (Skaletsky et al. 2003) and considered loci with confirmed transcription but without significant ORFs to be non-coding (Table S3, Data S3 & S4).

### Gene expression analyses

We measured transcript levels across adult SHR rat tissues using our own testis data and publicly available mRNA sequencing data for eight adult somatic tissues (Table S5) (Chang et al. 2023; Pattee et al. 2022; Schafer et al. 2015; Senko et al. 2022; van Heesch et al. 2019; Witte et al. 2021; Yang et al. 2020; Zeng et al. 2023). Within the testis, we used published data from five cell fractions from the rat testis (Table S5) (Chalmel et al. 2014; Fice and Robaire 2023; Wang et al. 2022).

We built a custom transcriptome by combining the transcripts from our annotation of sex chromosomes with X and autosomal transcripts from Ensembl release 104 (Dyer et al. 2025). For all datasets, we mapped RNA-seq reads to our custom transcriptome using kallisto with sequence-based bias correction (Bray et al. 2016) to estimate gene expression levels in transcripts per million (TPM). For multicopy gene families, we summed the number of reads that mapped to any single member of the gene family.

### Dot plots

We generated triangular dot plots (representing intrachromosomal sequence similarity) and square dot plots (representing interchromosomal sequence similarity) using a custom Perl script, available at http://pagelab.wi.mit.edu/material-request.html.

### Phylogenetic analyses

We aligned sequences with clustalw (version 2.1) (Larkin et al. 2007), and constructed trees using dnaml (version 3.66) from the Phylip package, using default parameters (Felsenstein 1989).

## Data Access

All BAC sequences generated in this study have been submitted to the NCBI Nucleotide Database (https://www.ncbi.nlm.nih.gov/nucleotide/), and accession numbers are listed in Supplemental Table S1. Whole genome sequencing data and RNA-seq data generated in this study have been submitted to the NCBI Sequence Read Archive (http://www.ncbi.nlm.nih.gov/sra) under accession numbers SRR32833872, SRR32834695, SRR1001913, and SRR26701686 as part of bioprojects PRJNA222509, PRJNA221163, and PRJNA226588. Supplemental material is available online at: http://pagelabsupplement.wi.mit.edu/papers/Bellott_et_al_2025/.

## Competing Interests

The authors declare no competing interests.

## Acknowledgments

This work was supported by the National Institutes of Health (R01HG007852 and U01HG007587), Whitehead Institute, the Howard Hughes Medical Institute, the Simons Foundation Autism Research Initiative, and generous gifts from The Brit Jepson d’Arbeloff Center on Women’s Health, Arthur W. and Carol Tobin Brill, Matthew and Hillary Brill, Charles Ellis, Carla Knobloch, the Brett Barakett Foundation, the Howard P. Colhoun Family Foundation, and the Seedlings Foundation.

## Author Contributions

Conceptualization: D.W.B., J.F.H., H.S., and D.C.P.; investigation: D.W.B., L.G.B., T.P., T.J.C., N.K., S.Z., D.K., M.M., S.D., Z.K., Q.W., J.W., D.M.M.; data curation, formal analysis, software, and validation: D.W.B., H.S., and J.F.H.; methodology, project administration: D.W.B., H.S., J.F.H., T.R., K.C.W., S.S., and D.M.M.; funding acquisition, supervision: T.R., R.A.G., and D.C.P.; visualization and writing – original draft: D.W.B.; writing – review & editing: D.W.B., J.F.H., H.S., and D.C.P.

## Notes

http://pagelabsupplement.wi.mit.edu/papers/Bellott_et_al_2025/

https://www.ncbi.nlm.nih.gov/bioproject/PRJNA222509

https://www.ncbi.nlm.nih.gov/bioproject/PRJNA221163

https://www.ncbi.nlm.nih.gov/bioproject/PRJNA226588

### Competing Interest Statement

The authors have declared no competing interest.

