## Supplemental_Figures for "The SHR/Akr Y chromosome reveals repeated turnover of the rat pseudoautosomal region"

Bellott, et al. 2025

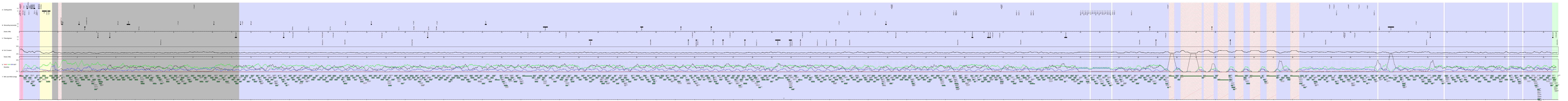
a Coding genes
b Noncoding transcripts
Scale (1Mb)
c Pseudogenes
d G+C Content
Scale (1Mb)
e SINE/LINE/ERV/SAT Densities
f BAC and WGS contigs

**Figure S1: Annotated sequence of SHR/Akr rat Y.**

Background colors represent sequence classes: pseudoautosomal (green); ancestral single-copy (yellow); X-transposed (red) ampliconic (blue); other (gray); heterochromatic (pink stripes). Gaps shown in white. All sequence features and BACs drawn to scale. Positions of all actively transcribed (a) intact protein-coding genes; (b) non-coding transcripts; and (c) pseudogenes. Plus (+) strand above, minus (-) strand below. (d) G+C content (%) calculated in a 100-kb sliding window with 1-kb steps. (e) SINE (red), LINE (green), ERV (blue), and Satellite (black) repeat densities, calculated in a 200-kb sliding window with 1-kb steps. (f) Sequenced BACs and WGS contigs. Each bar represents the size and position of a sequence within the assembly. Sequences with no prefix, RNAEX library; prefix "E", RNECO library; prefix "Link" WGS contigs supported by nanopore reads. Dark green, sequences in the tiling path; dark grey bars, other finished BAC clones; light grey, unfinished BAC clones.

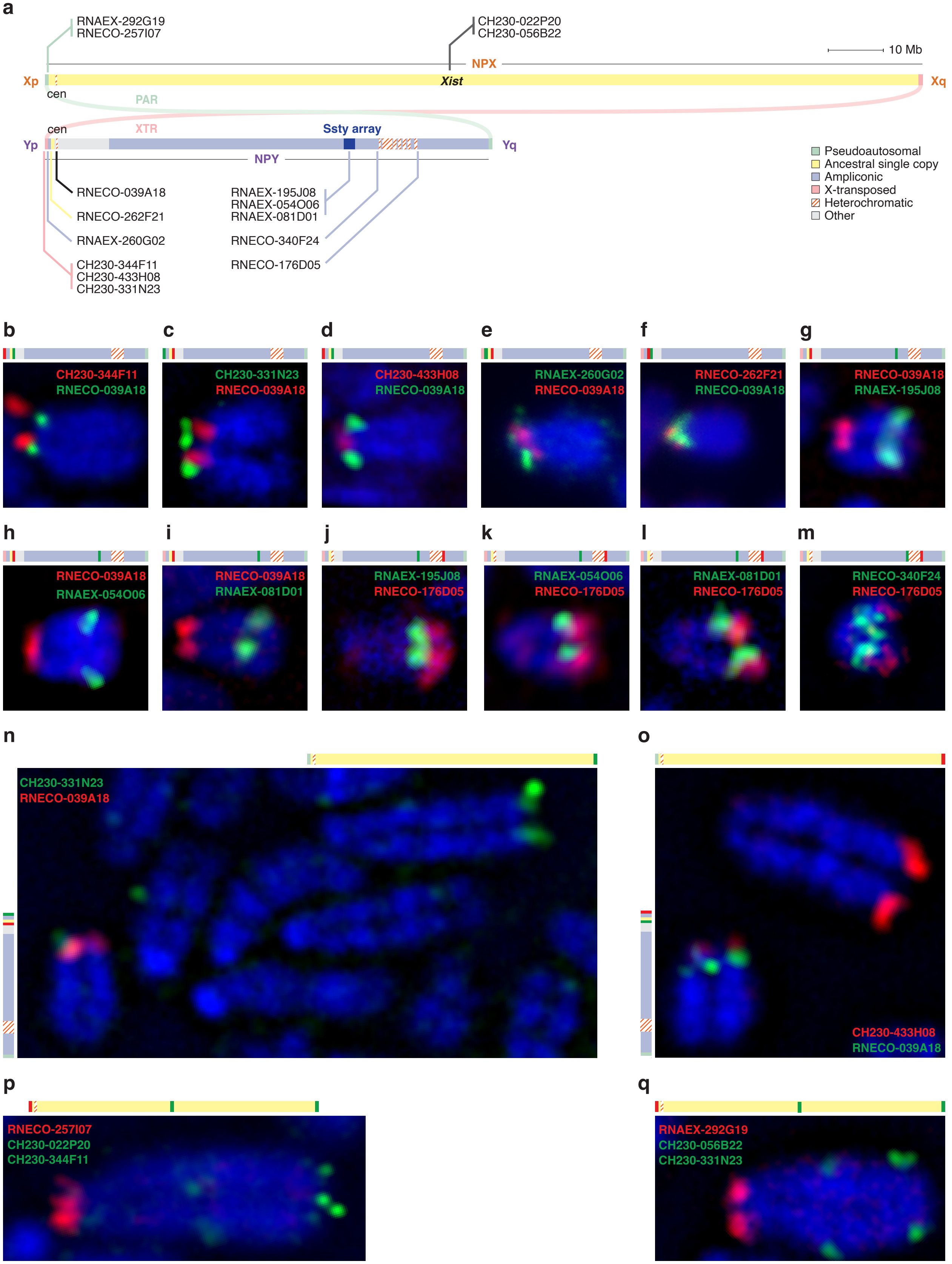

**Figure S2: Fluoresence in situ hybridization mapping of the rat sex chromosomes.**
(a) Schematic representation of the rat sex chromosomes indicating positions of BAC probes. Shading indicates six sequence classes. Scale bar 10 Mb. (b-q) Metaphase FISH with BAC probes with schematics indicating probe location on the rat sex chromosomes. (b-f) Short and (g-m) long arm probes on the rat Y. (n-o) XTR probes on the short arm of the Y and long arm of the X. (p-q) PAR probes (red) on the short arm of the X, with probes for *Xist* and the XTR (green) on the long arm.

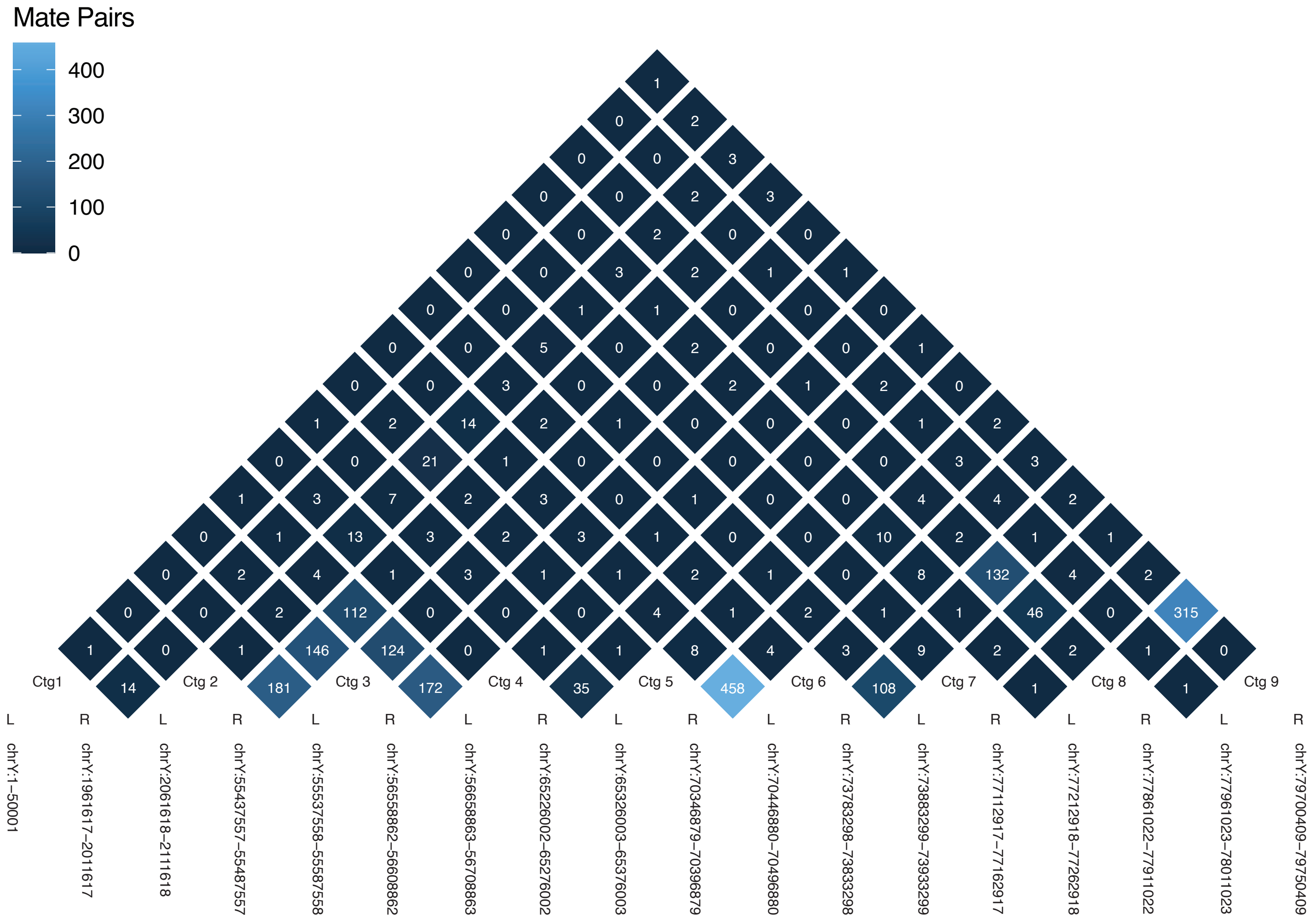

**Figure S3: HiC mapping of the rat Y chromosome.**
Heat map indiciating the number of HiC mate pairs shared between 50kb flanks of sequence contigs on the rat Y. Both reads in each mate pair uniquely map to sequence containing 32-mers that are unique in the rat genome.

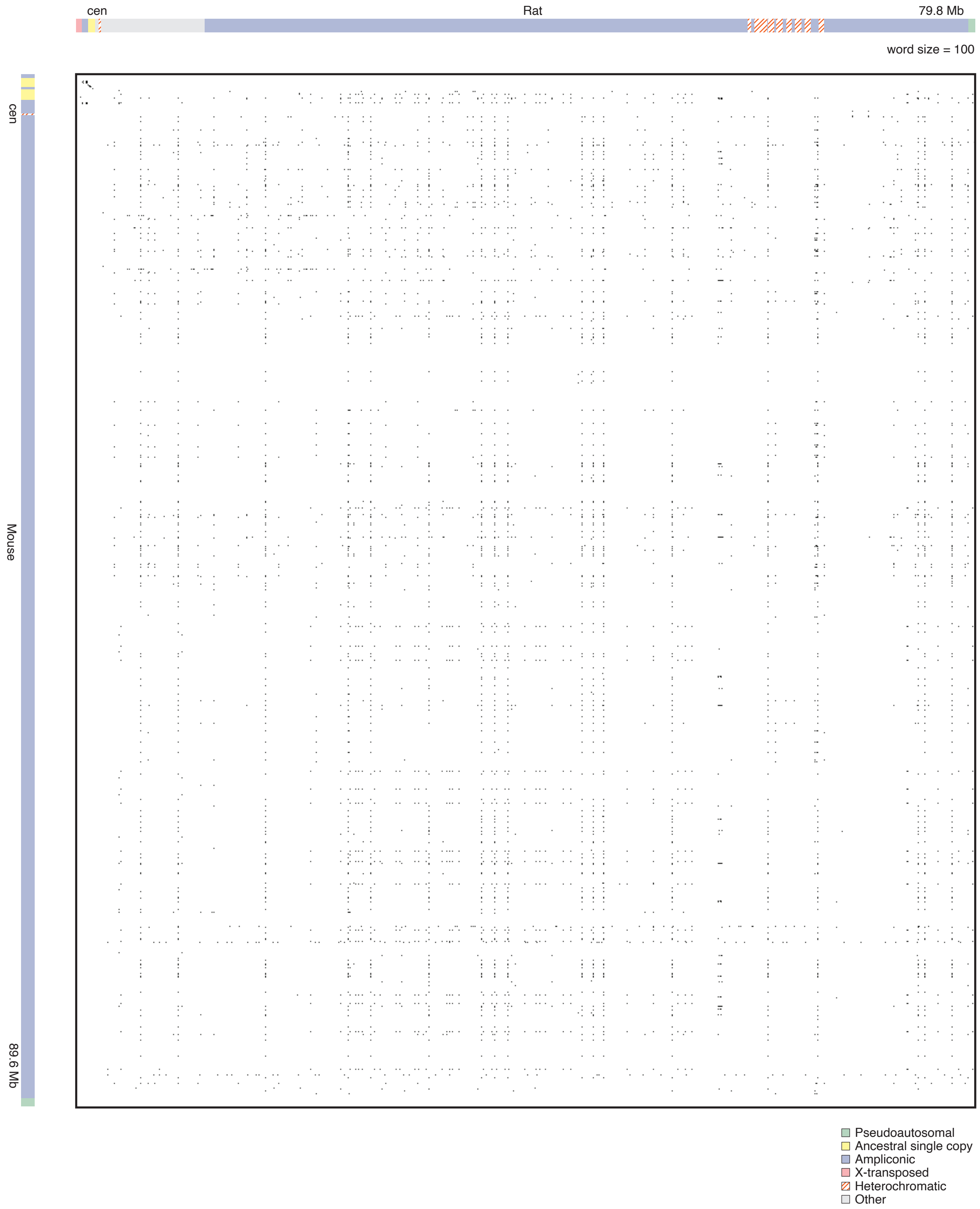

**Figure S4: Dot plot comparing rat and mouse Y chromosomes.**
Rectangular dot plot comparing the rat and mouse Y chromosomes. Each dot represents 100% identity in a 100-bp window. Schematic representation of the rat and mouse Y chromosomes shaded to indicate six sequence classes.

**Figure S5: Maximum-likelihood tree of *Ssty* family members in rat.** Maximum-likelihood tree of *Ssty* family members and spindlin family homologs in rat; genes shaded by chromosomal location: dark purple, NPY; light purple, NPY pseudogenes; orange, NPX; grey, autosomal pseudogenes. Heavy lines indicate branches leading to copies with intact ORFs; numbers, branches with greater than 50% bootstrap support. Scale bar; branch length in substitutions per site.

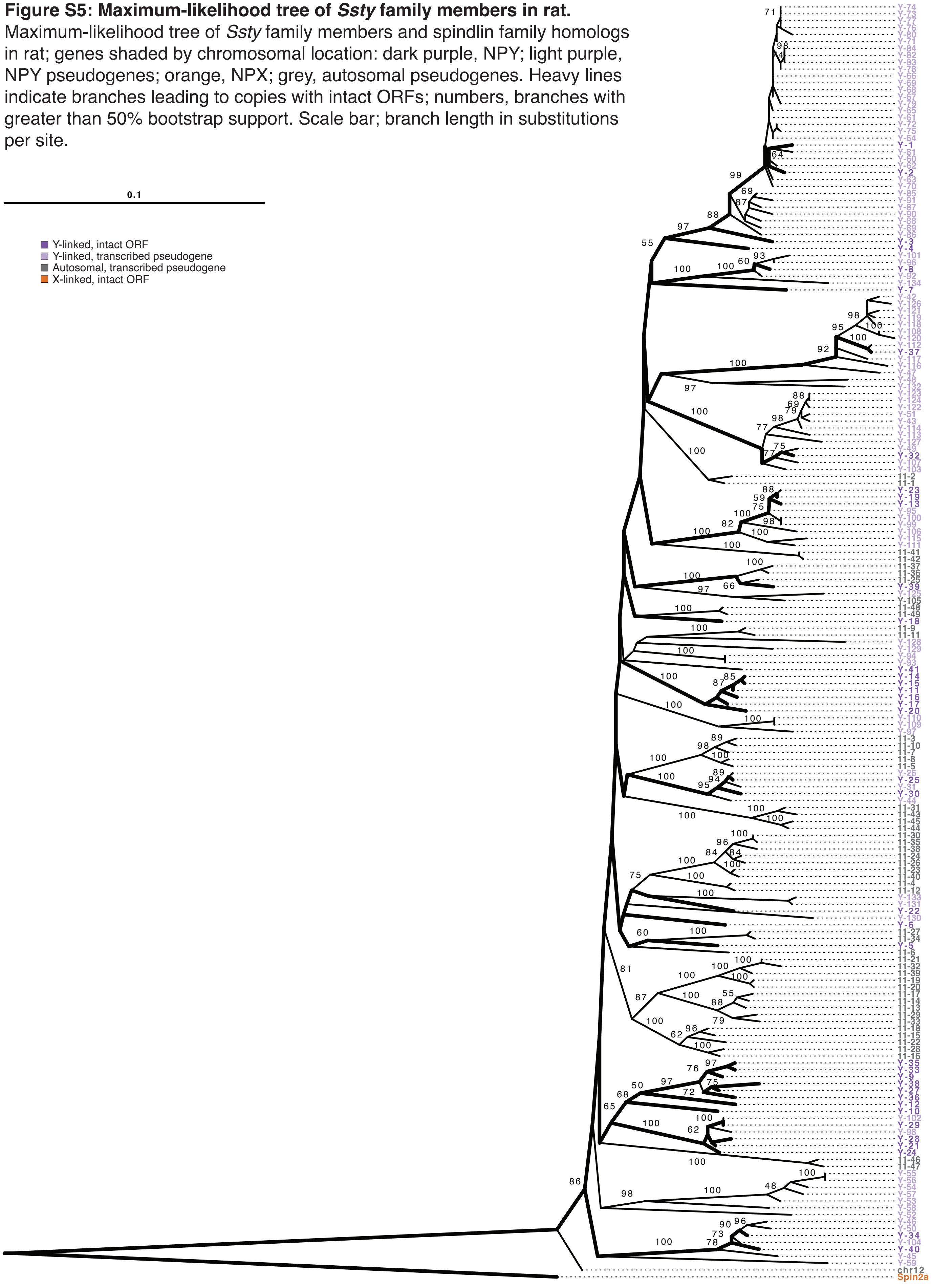

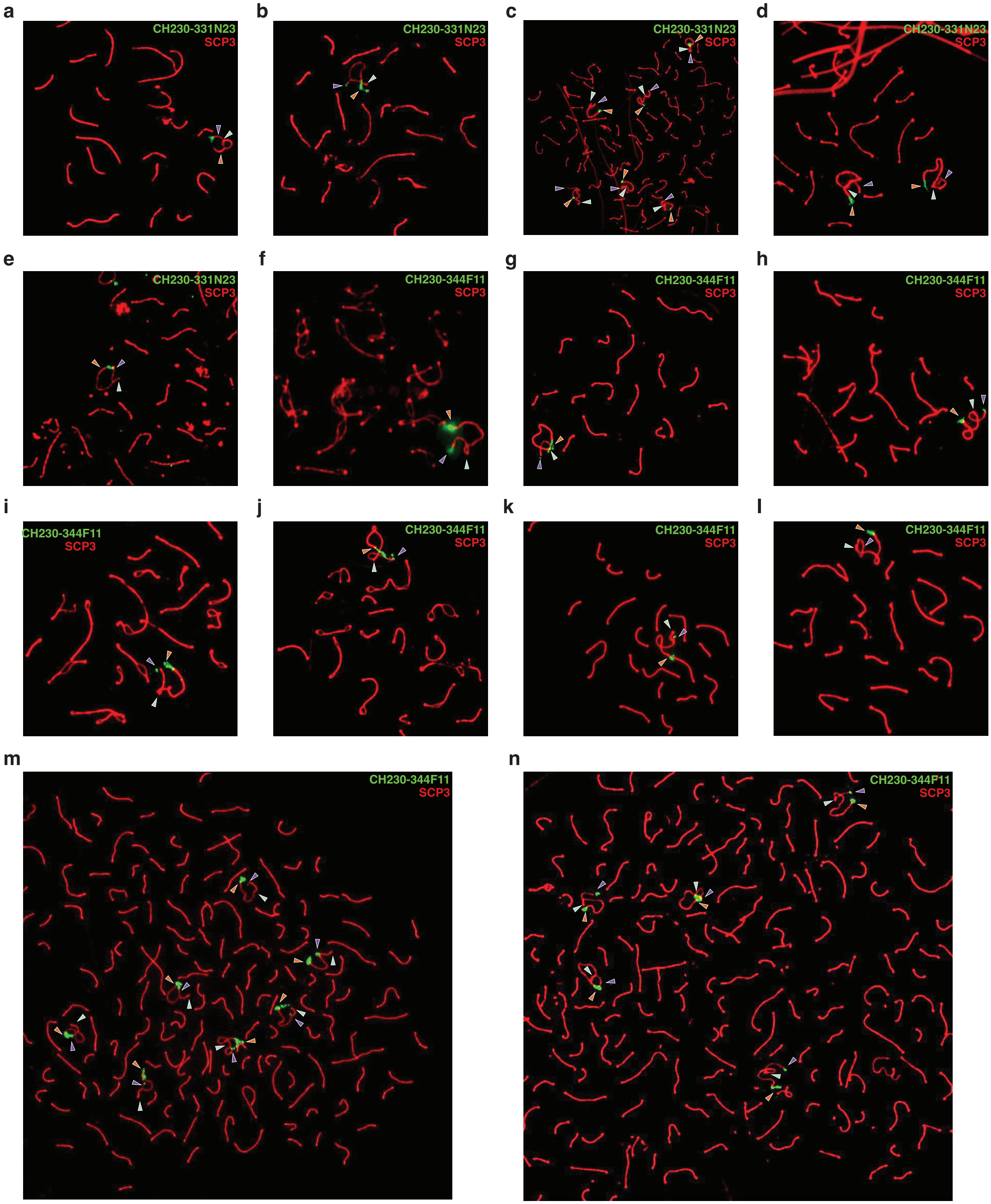

**Figure S6: Additional meiotic spreads showing location of X-transposed region BAC clones.**
FISH with X-transposed region BAC probes (a-e) CH230-331N23 and (f-n) CH230-344F11 (green) on meiotic prophase chromosomes, and anti-SCP3 antibody (red). Purple arrowhead, chrY; orange arrowhead, chrX; green arrowhead, PAR.
